# Self-Administration of entactogen psychostimulants dysregulates GABA and Kappa Opioid Receptor signaling in the central nucleus of the amygdala of female Wistar rats

**DOI:** 10.1101/2021.09.24.461477

**Authors:** Sophia Khom, Jacques D. Nguyen, Sophia A. Vandewater, Yanabel Grant, Marisa Roberto, Michael A. Taffe

## Abstract

Male rats escalate intravenous self-administration of entactogen psychostimulants, 3,4-methylenedioxymethcathinone (methylone) and 3,4-methylenedioxymethamphetamine (MDMA) under extended access conditions, as with typical psychostimulants. Here, we investigated whether female rats escalate self-administration of methylone, 3,4-methylenedioxypentedrone (pentylone), and MDMA and then studied consequences of MDMA and pentylone self-administration on GABA_A_ receptor and kappa opioid receptor (KOR) signaling in the central nucleus of the amygdala (CeA), a brain area critically dysregulated by extended access self-administration of alcohol or cocaine. Adult female Wistar rats were trained to self-administer methylone, pentylone, MDMA (0.5 mg/kg/infusion), or saline-vehicle using a fixed-ratio 1 response contingency in 6-hour sessions (long-access: LgA) followed by progressive ratio (PR) dose-response testing. The effects of pentylone-LgA, MDMA-LgA and saline on basal GABAergic transmission (miniature postsynaptic inhibitory currents, mIPSCs) and the modulatory role of KOR at CeA GABAergic synapses were determined in acute brain slices using whole-cell patch-clamp. Methylone-LgA and pentylone-LgA rats similarly escalated their drug intake (both obtained more infusions compared to MDMA-LgA rats) however, pentylone-LgA rats reached higher breakpoints in PR tests. At the cellular level, baseline CeA GABA transmission was markedly elevated in pentylone-LgA and MDMA-LgA rats compared to saline-vehicle. Specifically, pentylone-LgA was associated with increased CeA mIPSC frequency (GABA release) and amplitude (postsynaptic GABAA receptor function), while mIPSC amplitudes (but not frequency) was larger in MDMA-LgA rats compared to saline rats. In addition, pentylone-LgA and MDMA-LgA profoundly disrupted CeA KOR signaling such as both KOR agonism (1mM U50488) and KOR antagonism (200nM nor-binaltorphimine) decreased mIPSC frequency suggesting recruitment of non-canonical KOR signaling pathways. This study confirms escalated self-administration of entactogen psychostimulants under LgA conditions in female rats which is accompanied by increased CeA GABAergic inhibition and altered KOR signaling. Collectively, our study suggests that CeA GABA and KOR mechanisms play a critical role in entactogen self-administration like those observed with escalation of alcohol or cocaine self-administration.

## Introduction

The entactogen psychostimulant drugs 3,4-methylenedioxymethamphetamine (MDMA), 3,4-methylenedioxymethcathinone (Methylone) and 3,4-methylenedioxypentedrone (Pentylone) are commonly abused substances. MDMA, Methylone and Pentylone are monoamine transporter inhibitors and substrates with increased selectivity for serotonin over dopamine or norepinephrine transporters (Baumann et al., 2011;Simmler et al., 2013;Simmler et al., 2014a). Importantly, MDMA, Methylone and Pentylone are structurally closely related such that MDMA differs from Methylone by only the ketone on the beta carbon, while Methylone differs from Pentylone with respect to the length of the α-alkyl chain (Simmler et al., 2014b). Previous intravenous self-administration (IVSA) studies in male rats indicated that MDMA exhibits low efficacy as a reinforcer, leading to low overall drug intake and high inter-subject variability compared with, e.g., cocaine or methamphetamine (Bradbury et al., 2014; Creehan et al., 2015; Dalley et al., 2007).This has long been assumed to be a consequence of the pharmacological selectivity of MDMA for serotonin transporter inhibition and efflux, compared with the closely-related methamphetamine. However, previous studies demonstrated that MDMA is a more effective reinforcer when animals are initially trained to self-administer mephedrone (Creehan et al., 2015), or under higher ambient temperature conditions (Aarde et al., 2017; Cornish et al., 2008, 2003). Furthermore, male rats will obtain more infusions of MDMA when trained under daily extended or long-access sessions (6-hour) compared to short access (2-hour) sessions (Vandewater et al., 2015). Finally, despite 4-methylmethcathinone exhibiting preferential serotonin release (Kehr et al., 2011; Wright et al., 2012), similar to MDMA, it is a robust reinforcer in rat IVSA models (Creehan et al., 2015; Hadlock et al., 2011; Marusich et al., 2021; Nguyen et al., 2017). Thus there is evidence that under some circumstances, the serotonin transporter selective entactogen class stimulants can produce compulsive drug seeking behavior in rodent IVSA.

Within the class of entactogen stimulants, the propensity to support robust self-administration may vary. Pentylone appears to be more efficacious as a reinforcer than Methylone in a dose-substitution comparison in male and female rats originally trained to self-administer methamphetamine and α-pyrrolidinopentiophenone, respectively (Dolan et al., 2018; Javadi-Paydar et al., 2018). This may be because Pentylone exhibits reduced efficacy as a monoamine transporter substrate compared to MDMA or Methylone (Dolan et al., 2018), and displays less serotonin selectivity as a monamine transporter inhibitor relative to MDMA (Baumann et al., 2012; Simmler et al., 2016; Linda D. Simmler et al., 2014). This pharmacological profile suggests that Pentylone would be a highly efficacious reinforcer in rat IVSA procedures but it has not been well characterized apart from the two above-mentioned dose substitutions studies in animals trained on other drugs. Importantly, there are only limited data available that elucidate the abuse liability of entactogen stimulants in female subjects. These data show that, at least under 2-hour access conditions, the IVSA of Mephedrone(4-methylmethcathinone), Methylone and MDMA do not differ dramatically between male and female rats (Creehan et al., 2015; Javadi-Paydar et al., 2018; Vandewater et al., 2015). While Methylone IVSA is similar to MDMA IVSA when male rats are permitted 2h daily sessions, Methylone appears to be much more effective than MDMA under 6h daily access conditions (Nguyen et al., 2017; Vandewater et al., 2015). Therefore, subtle differences in IVSA methods may either reveal or obscure differences in abuse liability. This may be critical for the accuracy of inferences made about two or more closely-related entactogen psychomotor stimulants.

Thus, one major goal of this study was to determine if long-access to IVSA of three entactogens leads to escalating drug intake in female rats, as it does in males. As has been reviewed, it is increasingly recognized as important to confirm similarities and differences that may obtain between the sexes in a range of biomedical and neuroscience investigations (Clayton and Collins, 2014; Shansky and Murphy, 2021). A second goal was to test the hypothesis that extended access sessions would lead to increased IVSA of methylone relative to MDMA, as predicted by the indirect comparison of male long-access IVSA data (Nguyen et al., 2017; Vandewater et al., 2015). Lastly, we aimed to investigate neuroadaptations in synaptic transmission in the central nucleus of the amygdala (CeA) given its key role in the acute reinforcing actions of drugs of abuse as well the negative emotional state associated with drug withdrawal (Koob and Volkow, 2016). The CeA is composed primarily of GABAergic neurons and represents the major output area of the larger amygdaloid complex (Gilpin et al., 2015; Roberto et al., 2020). Chronic administration of drugs of abuse including ethanol (Gilpin et al., 2015; Kirson et al., 2021; Roberto et al., 2010, 2004), cocaine (Kallupi et al., 2013; Schmeichel et al., 2017; Sun and Yuill, 2020), methamphetamine (Li et al., 2015) or opioids (Bajo et al., 2014, 2011; Kallupi et al., 2020) enhance CeA GABA transmission representing a key molecular mechanism underlying maladaptive behaviors associated with addiction. Importantly, the CeA expresses several pro- and anti-stress promoting systems regulating its neuronal activity including the dynorphin/kappa opioid receptor (KOR) system, and chronic administration of drugs of abuse recruits these CeA stress systems (Koob, 2021; Koob and Schulkin, 2019). Specifically, cocaine-LgA is associated with a profound recruitment of CeA dynorphin/KOR signaling such as blockade of CeA KOR signaling reduces anxiety-like behaviors and cocaine-induced locomotor sensitization. Interestingly, cocaine-LgA also lead to a profound dysregulation of the CeA dynorphin/KOR system at the molecular level such as the KOR agonist U50488 increased CeA GABA release while the KOR antagonist nor-binaltorphimine decreased it (Kallupi et al., 2013). However, it has not yet been investigated whether or how MDMA-LgA or Pentylone-LgA affect CeA neuronal activity including GABAergic transmission and its regulation by the dynorphin/KOR system.

Thus, here we used acquisition of self-administration under long-access (6-hour) conditions, and post-acquisition dose substitutions under a Progressive Ratio schedule of reinforcement to assess potential differences in behavioral patterns in entactogen self-administration. For example, steeper escalation during LgA acquisition, or upward shifts in dose-response functions, are often inferred to represent meaningful differences in “addictiveness”. However, a difference in training-dose can appear to show differential “escalation” of IVSA of the same drug, such as with methamphetamine (Kitamura et al., 2006), and animals trained on a more-efficacious drug will respond for more of a less-efficacious drug, compared with those trained on the latter (Creehan et al., 2015; Vandewater et al., 2015). To determine if similar neuroadaptations are produced by long-access self-administration of drugs which produced different behavioral patterns, we performed *ex vivo* slice electrophysiology to assess changes in CeA GABA transmission and its regulation by the dynorphin/KOR system in female MDMA-LgA and Pentylone-LgA rats.

## Materials and Methods

### Animals

Female (N=56) Wistar rats (Charles River, New York) entered the laboratory at 10 weeks of age and were housed in humidity and temperature-controlled (23±1°C) vivaria on 12:12 hour light:dark cycles. Animals had *ad libitum* access to food and water in their home cages. All experimental procedures took place in scotophase and were conducted under protocols approved by the Institutional Care and Use Committees of The Scripps Research Institute and in a manner consistent with the Guide for the Care and Use of Laboratory Animals (National Research Council (U.S.). Committee for the Update of the Guide for the Care and Use of Laboratory Animals. et al., 2011).

### Drugs

Pentylone·HCl and Methylone·HCl were obtained from Cayman Chemical. 3,4-methylenedioxymethamphetamine (MDMA) HCl was obtained from NIDA Drug Supply. The MDMA analog was obtained from Fox Chase Chemical Diversity Center (Doylestown, PA, USA). Drugs were dissolved in physiological saline for the i.v. routes of administration. Dosing is expressed as the salt. We purchased tetrodotoxin (TTX) from Biotium (Hayward, CA, USA), and AP-5, CGP55845A and DNQX, U-50488 and nor-binaltorphimine from Tocris (Bristol, UK) for the electrophysiological recordings. Stock solutions of the drugs were prepared in either distilled water or dimethyl sulfoxide (DMSO) and added to the bath solution to achieve the desired concentration.

### Intravenous catheterization

Rats were anesthetized with an isoflurane/oxygen vapor mixture (isoflurane 5 % induction, 1-3 % maintenance) and prepared with chronic intravenous catheters as described previously (Nguyen et al., 2017a;Nguyen et al., 2018). Briefly, the catheters consisted of a 14-cm length polyurethane-based tubing (MicroRenathane®, Braintree Scientific, Inc, Braintree MA, USA) fitted to a guide cannula (Plastics one, Roanoke, VA) curved at an angle and encased in dental cement anchored to an ∼3-cm circle of durable mesh. Catheter tubing was passed subcutaneously from the animal’s back to the right jugular vein. Catheter tubing was inserted into the vein and secured gently with suture thread. A liquid tissue adhesive was used to close the incisions (3M™ Vetbond™ Tissue Adhesive; 1469S B). A minimum of 4 days was allowed for surgical recovery prior to starting an experiment. For the first 3 days of the recovery period, an antibiotic (cephazolin) and an analgesic (flunixin) were administered daily. During testing and training, intravenous catheters were flushed with ∼0.2–0.3 ml heparinized (32.3 USP/ml) saline before sessions and ∼0.2–0.3 ml heparinized saline containing cefazolin (100 mg/ml) after sessions. Catheter patency was assessed once a week, beginning in the third week of training, via administration through the catheter of ∼0.2 ml (10 mg/ml) of the ultra-short-acting barbiturate anesthetic, Brevital sodium (1 % methohexital sodium; Eli Lilly, Indianapolis, IN). Animals with patent catheters exhibit prominent signs of anesthesia (pronounced loss of muscle tone) within 3 s after infusion. Animals that failed to display these signs were considered to have faulty catheters and were discontinued from the study. Data that were collected after the previous passing of the test were excluded from analysis.

### Self-administration Procedure

#### Experiment 1 Acquisition

Following recovery from catheter implantation, rats were trained to self-administer MDMA (0.5 mg/kg per infusion; N=14), methylone (0.5 mg/kg per infusion; N=12), pentylone (0.5 mg/kg per infusion; N=15), or saline vehicle (N=8) using a fixed-ratio 1 (FR1) response contingency in 6-hour sessions. One individual in the MDMA group, 2 individuals in the methylone group and 2 individuals in the Pentylone group were lost due to nonpatent catheters. One individual in the Pentylone group was lost due to the catheter being chewed off by the cage mate. Operant conditioning chambers (Med Associates; Med-PC IV software) enclosed in sound-attenuating cubicles were used for self-administration studies as previously described (Nguyen et al., 2018, 2017). A pump pulse calculated to clear non-drug saline through the catheter started the session to ensure the first reinforcer delivery was not diluted, and a single priming infusion was delivered non-contingently if no response was made in the first 30 minutes of the session. Acquisition training was conducted for 14-15 sessions depending on the group so only the first 14 sessions are analyzed for the comparison.

#### Progressive-ratio (PR) dose-response testing

Rats in active drug groups were next subjected to dose substitution with the respective training drug (0.125, 0.5, 1.0, 2.5 mg/kg/infusion), followed by dose substitution with methamphetamine (0.01, 0.05, 0.1, 0.5 mg/kg/infusion), in a randomized order under a Progressive Ratio (PR) response contingency. One individual in the Pentylone group was lost due to the catheter being chewed off by the cage mate. The saline group completed five sequential PR sessions but again, only vehicle was available. For the PR, the sequence of response ratios started with one response then progressed thru ratios determined by the following equation (rounded to the nearest integer): Response Ratio = 5e^(injection number * j) – 5 (Richardson and Roberts, 1996). The value of “j” was 0.2 and was chosen so as to observe a “breakpoint” within ∼3 hrs. The last ratio completed before the end of the session (1 h after the last response up to a maximum of 3 h sessions) was operationally defined as the breakpoint. Following assessment with the training drug, groups were permitted to self-administer methamphetamine doses (0.01, 0.05, 0.1, 0.5 mg/kg/infusion) in a randomized order under the same PR schedule of reinforcement.

#### Experiment 2 Acquisition

Following recovery from catheter implantation, rats were trained to self-administer MDMA (0.5 mg/kg per infusion; N=8), pentylone (0.5 mg/kg per infusion; N=11), or saline vehicle (N=4), using a fixed-ratio 1 (FR1). One individual in the MDMA group was euthanized for illness. Acquisition training was conducted for 11-14 sessions depending on the group so only the first 11 sessions are analyzed for the comparison. Following acquisition, rats were trained on a variable number of sessions (X-Y total including acquisition) awaiting euthanasia for electrophysiological recordings.

### Animals for electrophysiology

Electrophysiological recordings were performed from a total of 28 randomly chosen rats. Specifically, we recorded from 8 rats from the saline-control group, 14 rats from the MDMA-LgA group, and 6 rats from the Pentylone-LgA group. Tissue for electrophysiology was collected 18 hours after the last self-administration session at the time animals would anticipate the next self-administration session. Importantly, rats were allowed to freely cycle during the self-administration process, and estrous cycle stage for each rat was determined upon sacrifice to evaluate its potential impact on CeA physiology. Estrous cycle was assessed based on cytological appearance of vaginal smear after euthanasia as described in (McLean et al., 2012). However, rats from both the saline and MDMA-LgA group were mainly in either pro-estrus or estrus, while Pentylone-LgA rats were either in estrus or diestrus. Thus, based on the unequal representation of estrous cycle stages in the different groups, data for GABA signaling and KOR pharmacology were pooled.

### Slice preparation and electrophysiological recordings

Preparation of acute brain slices containing the central nucleus of the amygdala (CeA) and electrophysiological recordings were performed as previously described (Khom et al., 2020a,b; Steinman et al., 2020; Suárez et al., 2019; Varodayan et al., 2018). Briefly, deeply anesthetized rats (3-5% isoflurane anesthesia) were quickly decapitated, and their brains placed in an ice-cold oxygenated high-sucrose cutting solution composed of 206 mM sucrose, 2.5 mM KCl, 0.5 mM CaCl_2_, 7 mM MgCl_2_, 1.2 mM NaH_2_PO_4_, 26 mM NaHCO_3_, 5 mM glucose, and 5 mM HEPES. We cut 300 µm thick coronal slices with the medial subdivision of the central amygdala (CeA) using a Leica VT 1000S and incubated them for 30 minutes in 37°C warm, oxygenated artificial cerebrospinal fluid (aCSF), composed of (in mM) 130 NaCl, 3.5 KCl, 2 CaCl_2_, 1.25 NaH_2_PO_4_, 1.5 MgSO_4_, 24 NaHCO_3_, and 10 glucose, followed by another 30 minutes incubation at room temperature. We identified CeA neurons with infrared differential interference contrast optics using a 40x water-immersion objective (Olympus BX51WI), and a CCD camera (EXi Aqua, QImaging). Using whole-cell patch technique, we recorded from 135 neurons pharmacologically isolated, action-potential independent miniature inhibitory postsynaptic currents (mIPSC) by adding the sodium-channel blocker tetrodotoxin (500nM, TTX), blockers of glutamate-mediated neurotransmission (6,7-dinitroquinoxaline-2,3-dione, 20µM (DNQX) and DL-2-amino-5-phosphonovalerate, 30µM (AP-5)), and the GABAB receptor antagonist CGP55845A (1µM) to the bath aCSF solution. All neurons were held -60mV. We performed recordings in a gap-free acquisition mode with a 10 kHz sampling rate and 10 kHz low-pass filtering using a MultiClamp700B amplifier, Digidata 1440A, and pClamp 10 software (MolecularDevices, San Jose, CA, USA). We pulled patch pipettes from borosilicate glass (3-5mΩ, King Precision) and filled them with a KCl-based internal solution composed of 145 mM KCl, 5mM EGTA, 5mM MgCl_2_, 10mM HEPES, 2mM Mg-ATP, and 0.2mM Na-GTP; pH was adjusted to 7.2-7.4 using 1N NaOH. We recorded only from neurons with an access resistance (R_a_) <15MΩ and/or with a R_a_ change<20% during the recording, as monitored by frequent 10mV pulses.

### Data analysis and Statistics

The number of infusions obtained in the IVSA experiments was analyzed by repeated measures *rmANOVA* with Sessions (acquisition only) or Dose as within-subjects factors. Significant main effects from the r*mANOVA* were further analyzed with post hoc multiple comparisons analysis using the *Tukey* procedure for multi-level, and the *Dunnett* procedure for two-level factors. Two missing data points (caused by program failure) in the Pentylone-trained rats during Session 11 were interpolated from the values before and after the last Session.

Frequencies, amplitudes, and current kinetics including current rise and decay times of mIPSCs were analyzed using MiniAnalysis software (Synaptosoft, Decatur, GA, USA). Data are given as means±S.E.M of raw values for mIPSC basal characteristics or from normalized values when assessing the effects of the KOR agonist U-50488 or the KOR antagonist nor-binaltorphimine (norBNI) on mIPSCs. Differences in mIPSC baseline characteristics were determined by a one-way *ANOVA* and a *Dunnett* post-hoc analysis. *Per se* effects of U-50488 or norBNI on mIPSCs were calculated by *one-sample t-tests*, and differences in drug effects across treatments was then also determined by *one-way ANOVA* with *Dunnett* post-hoc analyses. The criterion for significant results for both behavioral and electrophysiological data was set at *P* < 0.05 and all analyses were conducted using Prism 7 for Windows (v. 7.03; GraphPad Software, Inc, San Diego CA).

## Results

### Female Wistar rats escalate self-administration of entactogen psychostimulants under extended access (6-hour) conditions

The mean number of infusions obtained by rats trained on vehicle saline (N=8) decreased across sections, whereas infusions obtained by rats trained on pentylone, methylone or MDMA (for structural formulae, see **Fig. 1**) increased across the 14-session acquisition interval with the lowest mean drug-intake observed in the MDMA group and highest in the Pentylone group (**Fig. 2A**). Analysis of the saline, MDMA (N=13), Methylone (N=10) and Pentylone (N=12) groups confirmed a main effect of Session [*F* (13, 481) = 15.2; *P* < 0.0001], of Group [*F* (3, 37) = 3.324; *P* = 0.03] and of the interaction of factors [*F* (39, 481) = 2.368; *P* < 0.0001], on infusions obtained. The post hoc test confirmed that infusions were significantly increased compared to the first session in the Methylone (Sessions 8-14), Pentylone (Sessions 5-14) and MDMA groups (Sessions 9, 11-14); no significant differences in infusions were confirmed within the Vehicle trained group. Additionally, the Pentylone group was significantly different from Vehicle group during Sessions 5 and 10-14 and from the MDMA group during Sessions 5-6,13-14. The drug-lever responding (%) was significantly higher compared to responding Session 1 (**Fig. 2B**). The *ANOVA* confirmed a significant main effect of Session [*F* (13, 481) = 5.799; *P* < 0.0001] but not of Group [*F* (3, 37) = 1.649; *P* = 0.1948] or of the interaction of factors [*F* (39, 481)= 5.799; *P* = 0.4339]. The post hoc test confirmed that drug-lever responding during Sessions 5-14 was significantly different from the first session, collapsed across groups. During the final 5 sessions of acquisition, Pentylone and Methylone groups exhibited >80% drug-associated lever responding. The MDMA group exhibited >80% drug-associated lever responding during the final 2 sessions.

**Figure 1.**
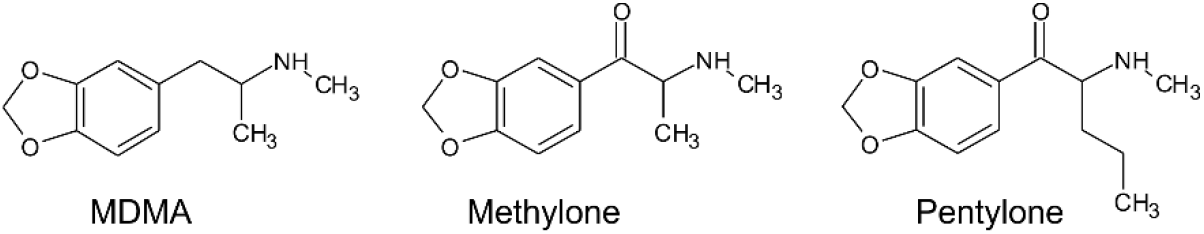
Structural formulae of MDMA, Methylone and Pentylone.

**Figure 2.**
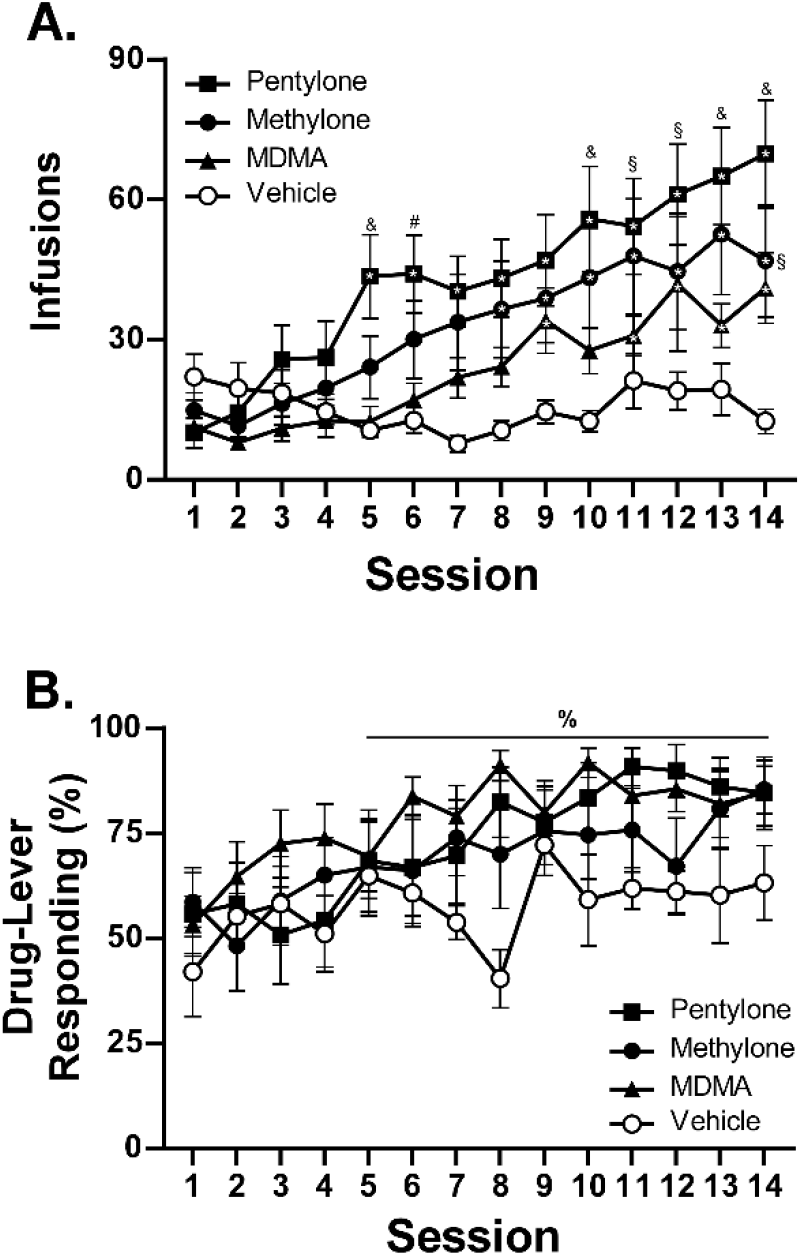
**A**) Mean (± S.E.M.) infusions of Pentylone (N=12), Methylone (N=10), MDMA(N=13), and saline (N=8; Vehicle) obtained under extended access conditions. **B**) Mean (± S.E.M.) percent of responses on the drug-associated lever. A significant difference from the first session, within group, is indicated with *, a significant difference from the first session, collapsed across groups, with %, a difference from the Vehicle and MDMA groups with &, a difference from the Vehicle group with §, and a difference from the MDMA group with #.

### Dose substitution in female Wistar rats following escalation of self-administration of entactogen psychostimulants

The rats trained on Pentylone (N=9), Methylone (N=7) or MDMA (N=10) under long-access conditions exhibited group differences during dose substitution experiments (**Fig. 3A**). Analysis confirmed a main effect of Dose [*F* (4, 92) = 30.04; *P* < 0.0001], of Drug [*F* (2, 23) = 6.067; *P* = 0.0077] and of the interaction of factors [*F* (8, 92) = 2.57; *P* < 0.05], on infusions obtained. Overall, rats trained on Methylone and Pentylone increased their intake to an approximately similar extent and received higher number of infusions compared to rats trained on MDMA. Pentylone-trained rats reached higher breakpoints than Methylone and MDMA-trained groups in PR tests.

**Figure 3.**
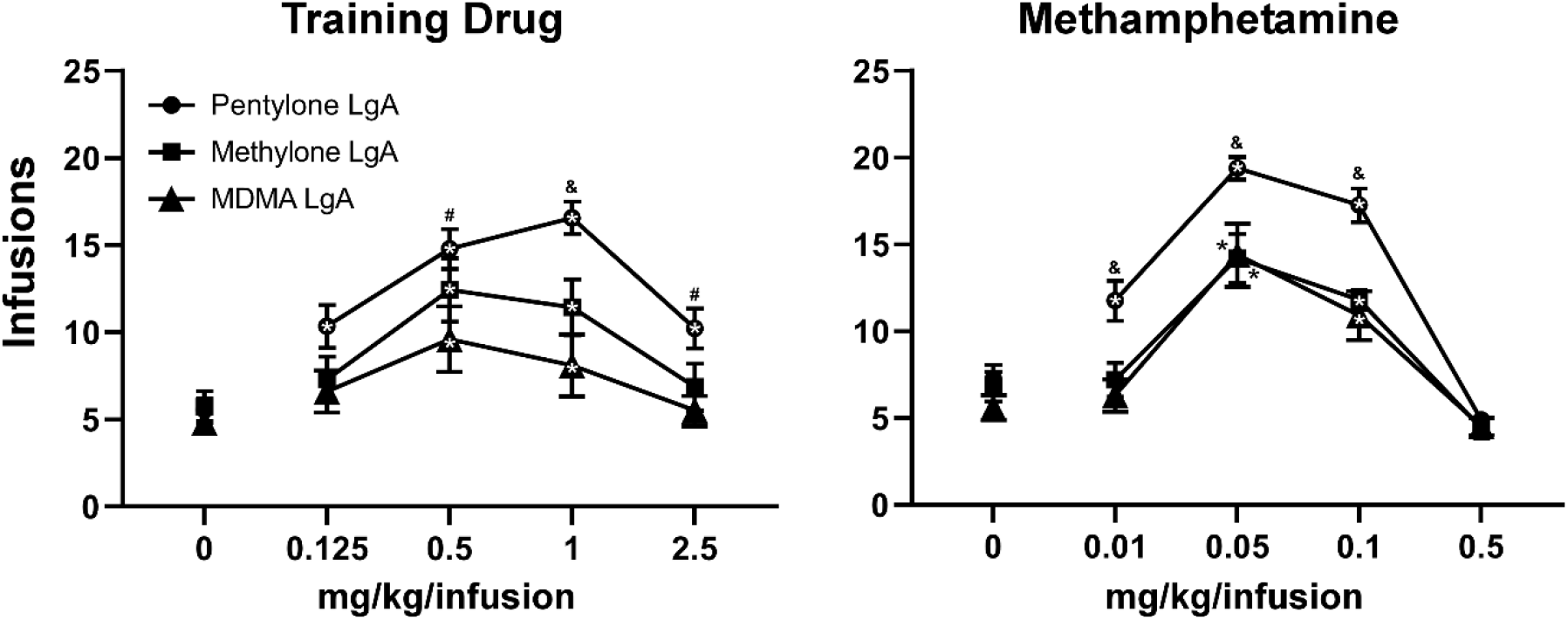
Mean (± S.E.M.) infusions of the respective training drug and of methamphetamine obtained by groups trained in LgA-IVSA of pentylone (N=8-9), methylone (N=5-7) or MDMA (N=10) are illustrated. A significant difference from saline, within group, is indicated with *, a significant difference from both other groups with &, and a difference from the MDMA LgA group with #.

When presented with methamphetamine substitution (**Fig. 3B**), Pentylone-LgA rats (N=8) similarly received higher number of infusions compared to rats trained on both Methylone-LgA (N=5) or MDMA-LgA rats. Analysis confirmed a main effect of Dose [*F* (4, 92) = 30.04; *P* < 0.0001], of Drug [*F* (2, 23) = 6.067; *P* = 0.0077] and of the interaction of factors [*F* (8,92) = 2.57; *P* < 0.05], on infusions obtained. One Pentylone animal that maintained patency was eliminated for exhibiting no dose sensitivity in the MA challenge, and two Methylone animals were eliminated due to failed catheter patency.

To further explicate the role of drug training history, the Pentylone-trained group were evaluated on doses of MDMA and the MDMA-trained group on doses of Pentylone, using the PR procedure. Pentylone supported higher levels of responding than did MDMA regardless of the training drug (**Fig 4**). Analysis confirmed a main effect of Dose [*F* (4, 132) = 35.75; *P*<0.0001], of Drug [*F* (3, 33) =12.47; *P*<0.0001] and of the interaction of factors [*F* (12, 132) = 2.098; *P*<0.05], on infusions obtained. The post hoc test confirmed that Pentylone-trained rats obtained a significantly higher number of infusions of Pentylone (0.125-2.5 mg/kg/infusion) compared to vehicle, and MDMA-trained rats also obtained more infusions of Pentylone than of vehicle (0.5-2.5 mg/kg/infusion). Similarly, each group obtained significantly more MDMA infusions (0.5 mg/kg/infusion) compared with vehicle. Within each drug, the groups did not differ, and exhibited similar dose-effect functions.

**Figure 4.**
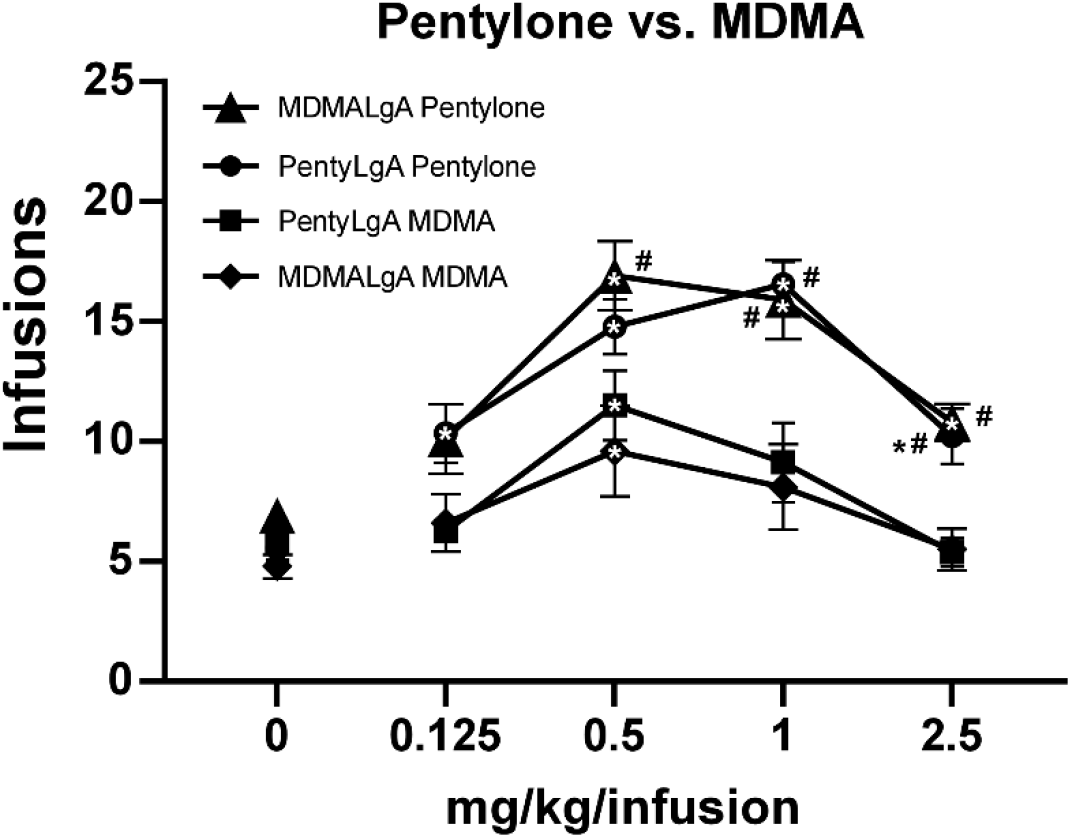
Mean (± S.E.M.) infusions of Pentylone and of MDMA obtained by groups trained in LgA IVSA of pentylone(N=8-9) or MDMA (N=10). A significant difference from saline, within group, is indicated with *, a significant difference from MDMA, within each LgA group, is indicated with #.

### Self-administration of MDMA or pentylone heightens CeA inhibitory signaling

Next, we assessed whether intravenous self-administration of MDMA (MDMA-LgA) or Pentylone (Pentylone-LgA) impacts CeA GABA transmission given that the CeA is highly sensitive to drugs of abuse such as alcohol or cocaine (Kallupi et al., 2013; Lesscher and Vanderschuren, 2012; Roberto et al., 2020; Schmeichel et al., 2017). We recorded pharmacologically isolated action-potential independent miniature inhibitory postsynaptic currents (mIPSCs) in 42 neurons from saline-control animals, 41 neurons from MDMA-LgA and 52 neurons from Pentylone-LgA rats. (Female rats that were selected for electrophysiological studies exhibited mean levels of drug-intake that were statistically indistinguishable from the rats that underwent behavioral testing only; **Fig. S1-3**). We found that MDMA-LgA and Pentylone-LgA increased CeA GABAergic transmission. Specifically, a *one-way ANOVA* (*F* (2. 132 = 5.021, *P* = 0.0079) with *Dunnett* post hoc analysis revealed that Pentylone-LgA but not MDMA-LgA significantly increased mIPSC frequencies compared to saline-controls (Saline: 0.74±0.09Hz vs. Pentylone-LgA: 1.12±0.09Hz, *P*= 0.0040 vs. MDMA-LgA: 0.90±0.08Hz, *P* = 0.3347)) suggesting enhanced vesicular GABA release (see **Fig. 5A, C**). Moreover, both pentylone-LgA and MDMA-LgA significantly increased mIPSC amplitudes (*one-way ANOVA*: *F* (2,132) = 7.617, *P* = 0.0007; *Dunnett* post hoc analysis: Saline: 57.1±2.2 pA vs. MDMA-LgA: 68.2±2.8pA, *P* = 0.0063 vs. Pentylone-LgA: 70.0±2.5pA, *P*= 0.0006, **Fig. 5B, D**) indicative of heightened postsynaptic GABA_A_ receptor function. Pentylone-LgA was further associated with faster mIPSC rise times (*one-way ANOVA*: *F* (2, 132) = 4.974, *P* = 0.0083) while mIPSC rise times were similar between MDMA-LgA and saline controls (*Dunnett* post hoc analysis: Saline: 2.67±0.04ms vs. MDMA-LgA: 2.61±0.04ms, *P* = 0.5142 vs. Pentylone-LgA: 2.50±0.04ms, *P* = 0.0051, **Fig. 5B, E**). Lastly, mIPSC decay times did not significantly differ between experimental groups (*F* (2,132) = 1.839, *P* = 0.1631, Saline: 9.0±0.3 ms vs. MDMA-LgA: 9.1±0.4ms vs. Pentylone-LgA: 8.3±0.3ms, see **Fig. 5B, F**). These data indicate that MDMA-LgA and Pentylone-LgA induce profound neuroadaptations to increase CeA GABA signaling which is a characteristic neuroadaptation observed after self-administration of other drugs of abuse.

**Figure 5.**
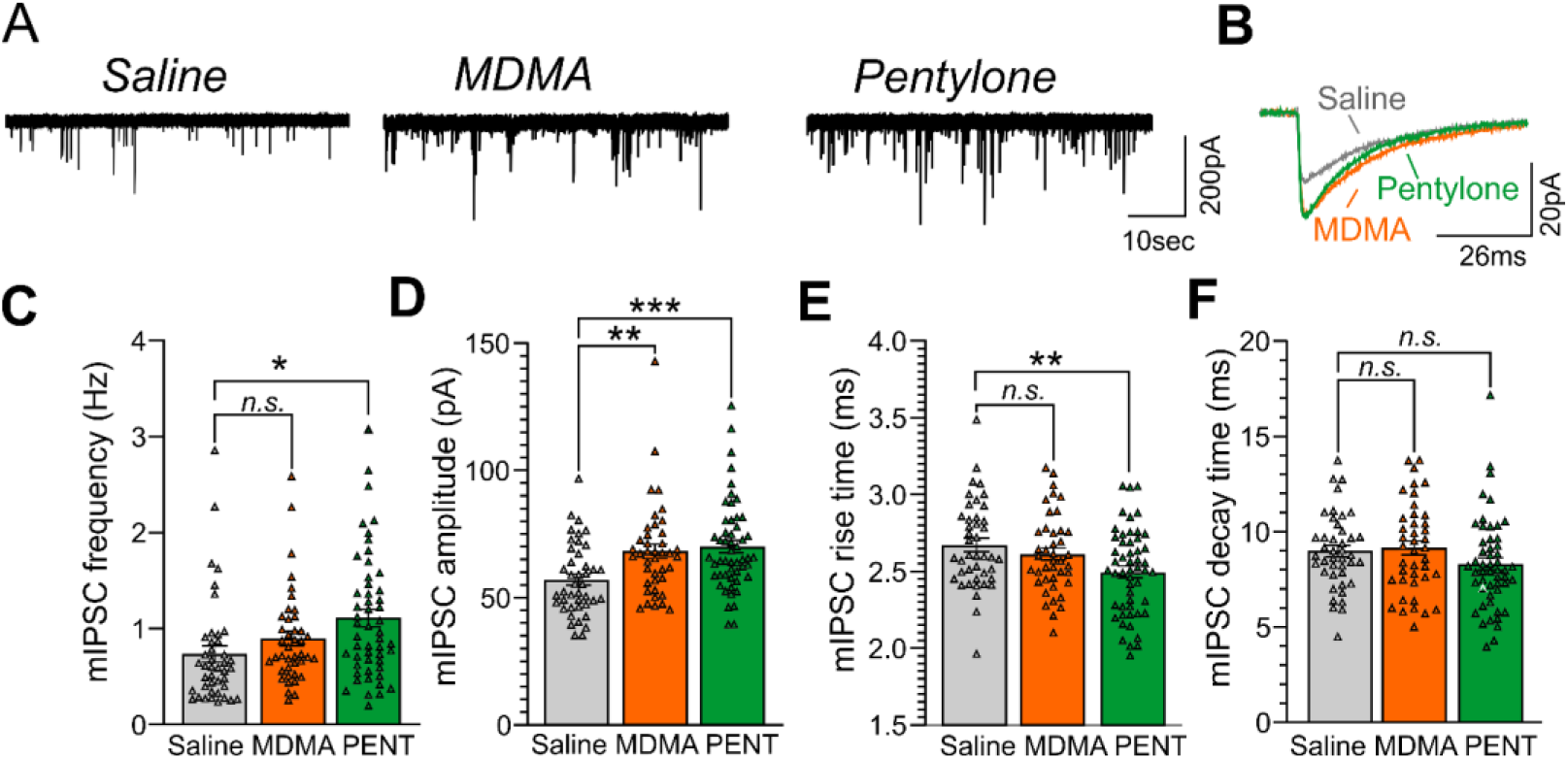
A) Representative mIPSC recordings from CeA neurons from female Wistar rats self-administering Saline (left panel), MDMA (middle panel), or Pentylone (abbreviated PENT, right panel). (B) Scaled mIPSC averages illustrating the effects of MDMA-LgA and Pentylone-LgA on mIPSC amplitudes and kinetics. Bars in represent means ± S.E.M. of mIPSC (C) frequencies, (D) amplitudes, (E) rise and (D) decay times. Differences between groups were calculated using a one-way ANOVA with Dunnet post hoc analyses. (*) = P < 0.05, (**) = P < 0.01, (***) = P < 0.001).

### MDMA and Pentylone self-administration disrupt endogenous KOR signaling

Given that CeA dynorphin/KOR signaling drives behaviors associated with excessive drug consumption including cocaine or alcohol self-administration (Anderson et al., 2019; Bloodgood et al., 2020; Kallupi et al., 2013; Koob, 2008), we lastly tested whether MDMA-LgA or Pentylone-LgA would also alter KOR-mediated regulation of vesicular CeA GABA release. As shown in **Fig. 6**, activating KOR by application of the selective agonist U-50488 (1µM as in (Gilpin et al., 2014; Kallupi et al., 2013)) in saline-controls significantly decreased mIPSC frequency (63.1±3.6%, *t* = 10.13, *df* =10, *P* < 0.0001, *one-sample t-test*) without affecting any postsynaptic measures indicating that KOR agonism reduces CeA presynaptic GABA release (**Fig. 6A, B**). Conversely, application of the KOR antagonist nor-binaltorphimine (norBNI, 200nM, as in (Gilpin et al., 2014; Kallupi et al., 2013)) increased mIPSC frequency (119.1±7.9%, *t* = 2.414, *df* = 9, *P* = 0.039) in saline-controls indicative of a tonic endogenous dynorphin/KOR signaling regulating GABA signaling under physiological conditions also in female rats (**Fig. 6A, C**). Moreover, norBNI did not alter postsynaptic properties of mIPSCs in saline-controls as has been previously reported (Kallupi et al., 2013).

**Figure 6.**
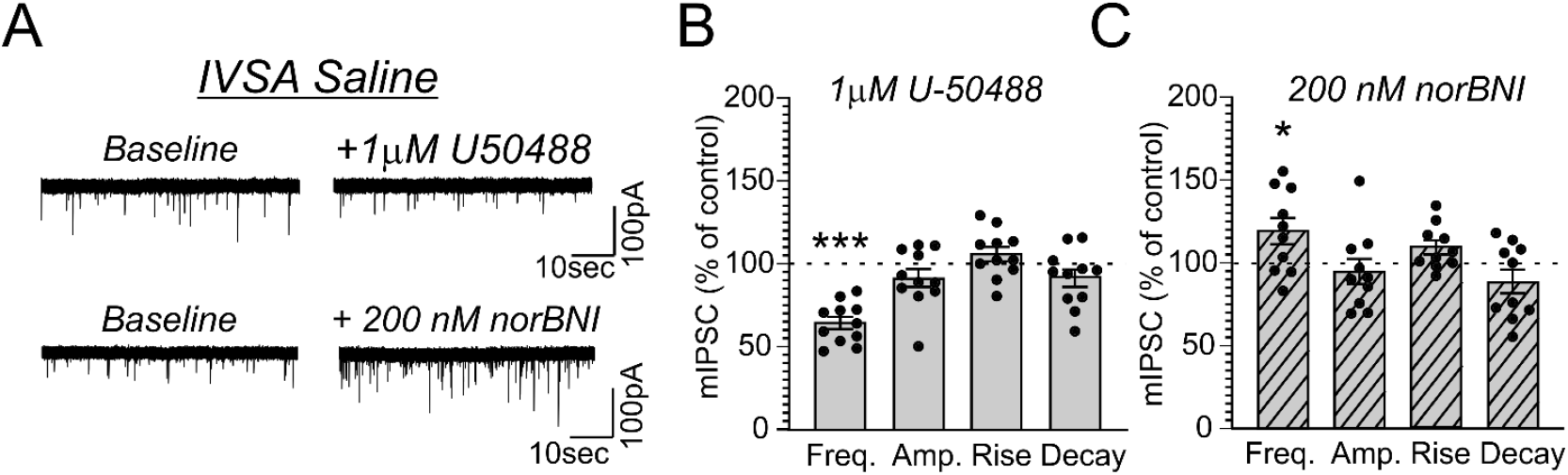
(A) Representative mIPSCs from CeA neurons during control and during superfusion with the KOR-agonist U-50488 (1µM, upper panel) or the KOR-antagonist norBNI (200nM, lower panel) are shown. Bars represent means ± S.E.M of the normalized effects either (B) U-50488 or (C) norBNI at the indicated concentration on mIPSC characteristics. Statistically significant differences to baseline control were calculated using a one-sample t-test. (*) = P < 0.05, (***) = P < 0.001.

Application of U-50488 (1µM) similarly decreased mIPSC frequency in MDMA-LgA (62.7±8.8%, *t* = 4.243, *df* = 10, *P* = 0.0017, *one-sample t-test*, **Fig.7A, B**) and Pentylone-LgA (50.5±6.4%, *t* = 7.747, *df* =11, *P* < 0.0001, *one-sample t-test*, **Fig. 8A, B**) rats. A *one-way ANOVA* analysis further confirmed that the effects of U-50455 on mIPSC frequency did not differ between saline, MDMA-LgA and Pentylone-LgA rats (*F* (2, 31) = 1.196, *P* = 0.3161). U-50488 did not alter mIPSC amplitudes and current kinetics in MDMA-LgA rats, but it significantly decreased mIPSC amplitudes in Pentylone-LgA rats without affecting mIPSC rise and decay times indicating that KOR-activation after Pentylone-self-administration also decreases postsynaptic GABA_A_ receptor function presumably leading to reduced neuronal inhibition. Moreover, a *one-way ANOVA* analysis confirmed highly significant differences in the effects of the KOR-antagonist norBNI on CeA vesicular GABA release in MDMA-LgA and Pentylone-LgA rats compared to saline-controls (*F* (2, 33) = 13.13, *P* < 0.0001). Specifically, unlike to the control group (Fig 6C) where we found that application of norBNI (200nM) increased mIPSC frequency, in both MDMA-LgA (79.0±7.5%, *t* = 2.682, *df* = 12, *P* = 0.02, *one-sample t-test*) and Pentylone-LgA rats (65.6±5.5%, *t* = 6.249, df = 11, *P* < 0.0001, *one-sample t-test*) norBNI decreased GABA release. Overall, this switch in the tonic role of KOR in modulating GABA release combined with the evidence that antagonist and agonist display a similar pharmacological profile suggests that both excessive MDMA and Pentylone self-administration under long-access conditions induce significant neuroadaptations of KOR receptor signaling. Lastly, norBNI did not significantly alter any postsynaptic mIPSC characteristics including amplitude, rise or decay times in either MDMA-LgA (**Fig.7C**) or Pentylone-LgA rats (**Fig. 8C**).

**Figure 7.**
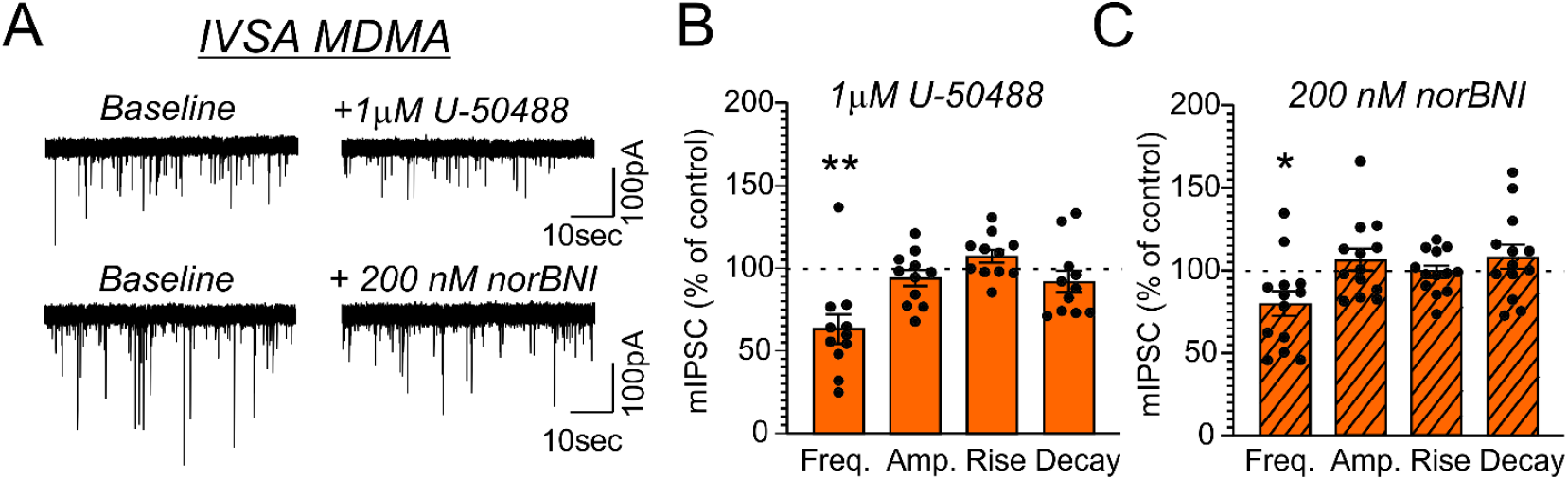
(A) Representative mIPSCs from CeA neurons during control and during superfusion with the KOR-agonist U-50488 (1µM, upper panel) or the KOR-antagonist norBNI (200nM, lower panel) are shown. Bars represent means ± S.E.M of the normalized effects of (B) U-50488 or (C) norBNI on mIPSC characteristics. Statistically significant differences to baseline control were calculated using a one-sample t-test. (*) = P < 0.05, (**) = P < 0.01.

**Figure 8.**
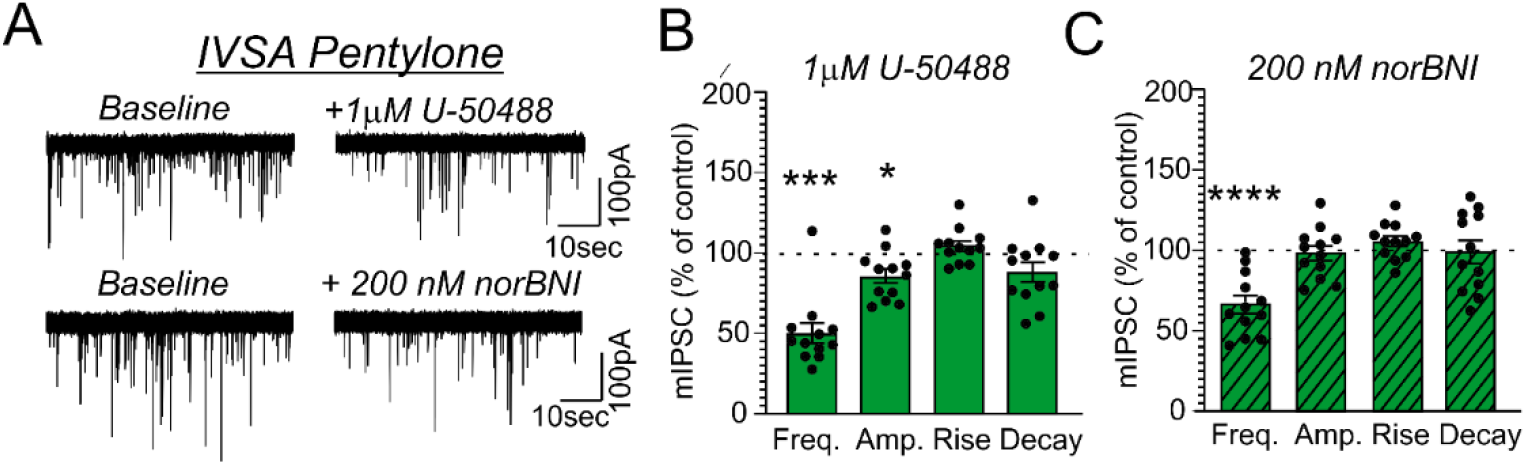
(A) Representative mIPSCs from CeA neurons during control and during superfusion with the KOR-agonist U-50488 (1µM, upper panel) or the KOR-antagonist norBNI (200nM, lower panel) are shown. Bars represent means ± S.E.M of the normalized effects either (B) U-50488 or (C) norBNI at the indicated concentration on mIPSC characteristics. Statistically significant differences to baseline control were calculated using a one-sample t-test. (*) = P < 0.05, (**) = P < 0.01.

## Discussion

This study shows that female rats readily acquire the self-administration of methylone, pentylone and MDMA under 6-hour long-access (LgA) daily training conditions. The groups trained on Methylone and Pentylone increased their intake to an approximately similar extent, with MDMA-trained animals increasing to a slightly lower extent, when considered as a population. This represents the first replication of extended-access IVSA intake of entactogen cathinones and MDMA that was previously reported for male rats trained to self-administer Methylone, Mephedrone or MDMA (Nguyen et al., 2017; Vandewater et al., 2015). Moreover, this is the first study to demonstrate a profound dysregulation of CeA neuronal activity in response to self-administration of entactogens in female rats. Together these results confirm that there is nothing qualitatively protective about the entactogens relative to other drugs of abuse, e.g., methamphetamine, cocaine or alcohol, and apparent differences in behavioral responding in intravenous self-administration procedures may be a function of the duration of action of a training dose (akin to what has been reported for methamphetamine; (Kitamura et al., 2006)). The post-acquisition dose-effect curves further emphasize that that in some cases the training history may (methamphetamine) or may not (Pentylone/MDMA) interact with the available drug to determine self-administration rate. However, systematic dose functions for highly effective reinforcers such as Pentylone and methamphetamine illustrated that all groups, regardless of training history, exhibited motivated drug-seeking behavior. One unexpected outcome was the self-administration of the MDMA analog, since it was constructed to be the amphetamine analog of Pentylone (**Fig. S4**). In the between groups analysis of the PR dose-substitution, Pentylone was more efficacious in comparison with Methylone (i.e., the cathinone analog of MDMA). The MDMA analog compound exhibited, if anything, reduced potency and similar efficacy relative to MDMA, represented by a rightward shift of the dose-response curve. This is a further caution against simplistic structure-activity inferences about *in vivo* activity in the intravenous self-administration procedure.

The drug substitution experiments show that inferences that MDMA is less addictive based on lower rates of intravenous self-administration during acquisition or in a dose-substitution procedure may be misleading. This was further confirmed by the electrophysiological experiments. The disruption of CeA synaptic transmission that has been associated with escalated self-administration of a range of drugs also occurred in the MDMA LgA group in this study. Effects were similar in entactogen trained groups that exhibited differences in behavioral drug intake. Specifically, we found that both MDMA-LgA and Pentylone-LgA exhibited markedly elevated CeA GABA transmission leading to enhanced local inhibition by either increasing presynaptic GABA release (Pentylone-LgA) and/or enhancing postsynaptic GABA_A_ receptor function (MDMA-LgA and Pentylone-LgA). Importantly, elevated inhibitory CeA signaling is a <u>key molecular mechanism</u> driving behaviors associated with drug abuse including escalation of drug intake in response to the emergence of the negative emotional state (Koob, 2021). Thus, our electrophysiological data indicate that Pentylone-LgA increased GABA transmission at both pre- and postsynaptic sites including potential changes in GABA_A_ receptor subunit composition leading to a presumably stronger CeA neuronal inhibition, while MDMA-LgA only elevated postsynaptic GABA_A_ receptor function but did not affect CeA GABA release.

Our study revealed that regulation of CeA synaptic GABA transmission by the dynorphin/KOR system in female rats does not differ from that in male rats (Gilpin et al., 2014; Kallupi et al., 2020); that is, activation of KOR in female rats also decreases CeA GABA release, while KOR antagonism increases CeA GABA transmission supporting a tonic role of KOR in the basal CeA GABA activity. Furthermore, both MDMA and Pentylone self-administration under long-access conditions disrupted CeA regulation by the dynorphin/KOR system. Specifically, we found that KOR activation with U-50488 decreased CeA GABA transmission in both MDMA-LgA and Pentylone-LgA rats mainly via reducing presynaptic GABA release. Moreover, the KOR antagonist norBNI did not *increase* CeA GABA transmission (as in saline controls) but *decreased* it in both MDMA-LgA and Pentylone-LgA rats. Interestingly, Pentylone-LgA animals escalated their drug intake significantly more than MDMA-LgA rats suggesting that distinct neuroadaptations within CeA GABAergic synapses, associated with more pronounced local inhibition, may potentially account for the observed differences in drug escalation. KOR activation with U-50488 decreased postsynaptic GABA_A_ receptor function only in Pentylone-LgA rats suggesting larger inhibitory effects of KOR activation on CeA GABAergic synapses after Pentylone-LgA.

Similar paradoxical effects of norBNI on CeA GABA signaling (i.e., norBNI decreasing CeA GABA transmission instead of increasing it) have been previously reported after cocaine-LgA (Kallupi et al., 2013). However, while after cocaine-LgA the effect of the KOR agonist on CeA GABA release had also changed directionality, i.e., KOR activation led to increased instead of decreased GABA signaling, in our study the KOR agonist U-50488 decreased CeA GABA release. This indicates some distinctions of the neuroadaptations at GABAergic synapses in response to cocaine-LgA vs. MDMA-LgA or Pentylone-LgA. Potentially, the fact that cocaine, MDMA and pentylone exhibit different mechanisms of action with respect to their activities at the different monoamine transporters (Baumann and Volkow, 2016; Glatfelter et al., 2021; Saha et al., 2019; Sandtner et al., 2016; Linda D. Simmler et al., 2014; Simmler et al., 2016; Steinkellner et al., 2011) may account for distinct neuroadaptations at CeA GABAergic synapses. Interestingly, the fact that norBNI did not increase CeA GABA release after Pentylone and MDMA self-administration may suggest a loss of tonic dynorphin signaling in the CeA at first sight, but it could also stem from alternative or non-canonical KOR signaling cascades resulting from repeated drug exposure. Indeed, KOR signaling has been shown to highly sensitive to stressful events and moreover, it induces activation of kinase cascades including G-protein coupled Receptor Kinases (GRK) and members of the mitogen-activated protein kinase (MAPK) family or β-arrestin-dependent pathways, amongst other classical G-protein mediated mechanisms (Bruchas and Chavkin, 2010; Ho et al., 2018; Lovell et al., 2015; Uprety et al., 2021). Thus, we hypothesize that the observed norBNI effects stem from changes in KOR signaling rather than a loss of CeA dynorphin, however, future studies utilizing different KOR antagonists will facilitate more insights into this phenomenon.

Overall, this study represents the first replication of extended-access IVSA intake of entactogen cathinones and MDMA in female rats, similar to that previously reported for male rats. The *in vivo* efficacy of cathinone compounds as reinforcers may not be supported by simplistic structure-activity inferences, nor by simplistic analysis of response rates in the acquisition of IVSA. Comparison of training groups across IVSA of the same compounds indicates a more similar motivational state. Furthermore, our studies also reveal similar profound neuroadaptations of CeA GABA transmission, and its regulation by the dynorphin/KOR system, in both MDMA-LgA and Pentylone-LgA groups, despite behavioral differences in the acquisition phase. Heightened GABA signaling associated with increased local inhibition in the CeA might represent a consistent, key mechanism underlying the escalation of drug self-administration.

## Author Contributions

This is manuscript number 30134 from the Scripps Research Institute. SK, JDN, MR and MAT designed the studies. SK, JDN, YG, and SAV performed the research and conducted initial data analysis. SK, JDN, MR and MAT conducted statistical analysis of data, created figures, and wrote the paper. All authors approved of the submitted version of the manuscript.

## Data Availability Statement

All data needed to evaluate the conclusions in this paper are present in the paper. Additional data related to this paper may be requested from the corresponding authors.

## Declaration of Transparency and Scientific Rigor

This paper adheres to the principles for transparent reporting and scientific rigor of preclinical research recommended by funding agencies, publishers and other organizations engaged with supporting research.

