## Supplement for "Self-Administration of entactogen psychostimulants dysregulates GABA and Kappa Opioid Receptor signaling in the central nucleus of the amygdala of female Wistar rats"

***Supplementary Information***

Table of Contents

Supplementary Results	2

Figure S1. Self-administration of Pentylone in Rats Selected for Behavioral-only or Electrophysiological Studies.	2

Figure S2. Self-administration of MDMA in Rats Selected for Behavioral-only or Electrophysiological Studies.	3

Figure S3. Self-administration of Saline in Rats Selected for Behavioral-only or Electrophysiological Studies.	4

Figure S4. Self-administration of MDMA or MDMA-analog in female rats trained to self-administer Pentylone.	5

### Supplementary Results

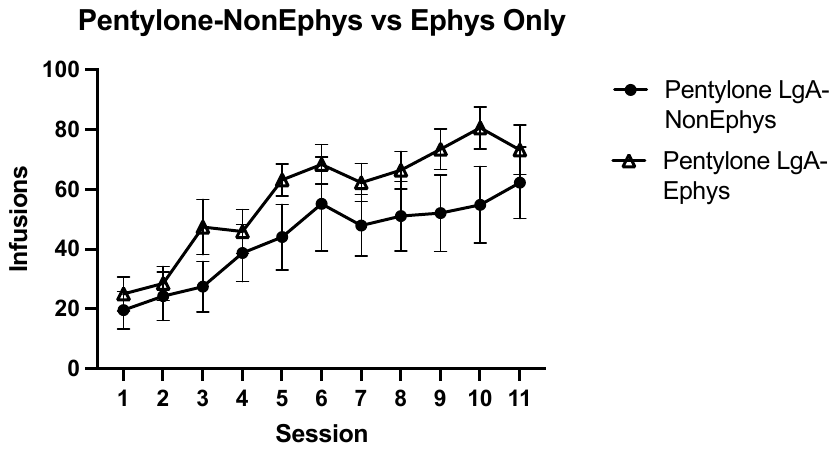

#### Figure S Self-administration of Pentylone in Rats Selected for Behavioral-only or Electrophysiological Studies.

*Mean (±SEM) pentylone infusions obtained by long-access rats (Pentylone LgA- NonEphys ; N=11) compared to long-access rats selected for ephysiological studies (Pentylone LgA- Ephys Only; N=14) across acquisition training sessions of self-administration. There was no significant difference in pentylone infusions between groups (P> 0.05). Significant difference from Session 1 within-group is indicated by #.*

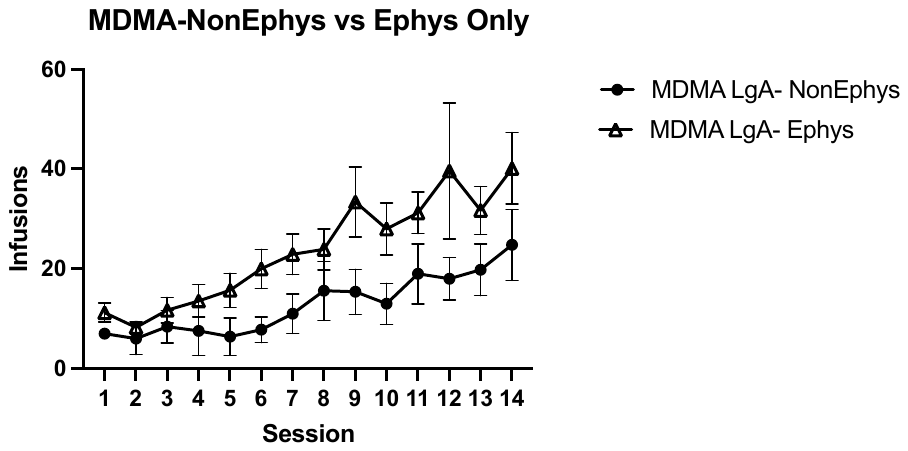

#### Figure S Self-administration of MDMA in Rats Selected for Behavioral-only or Electrophysiological Studies.

*Mean (±SEM) MDMA infusions obtained by long-access rats (MDMA LgA- NonEphys; N=5) compared to long-access rats selected for ephysiological studies (MDMA LgA- Ephys Only; N=14) across acquisition training sessions of self-administration. There was no significant difference in pentylone infusions between groups.*

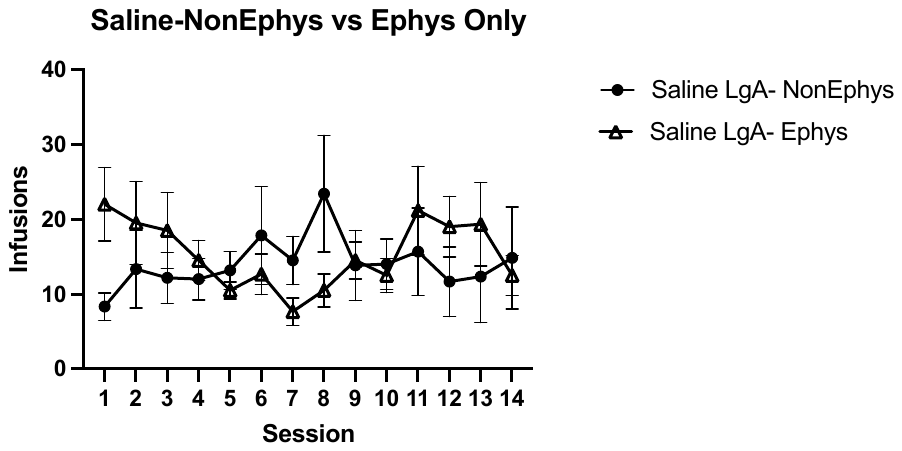

#### Figure S Self-administration of Saline in Rats Selected for Behavioral-only or Electrophysiological Studies.

*Mean (±SEM) saline infusions obtained by long-access rats (Saline LgA- NonEphys; N=6) compared to long-access rats selected for ephysiological studies (Saline LgA- Ephys Only; N=6) across acquisition training sessions of self-administration. There was no significant difference in pentylone infusions between groups.*

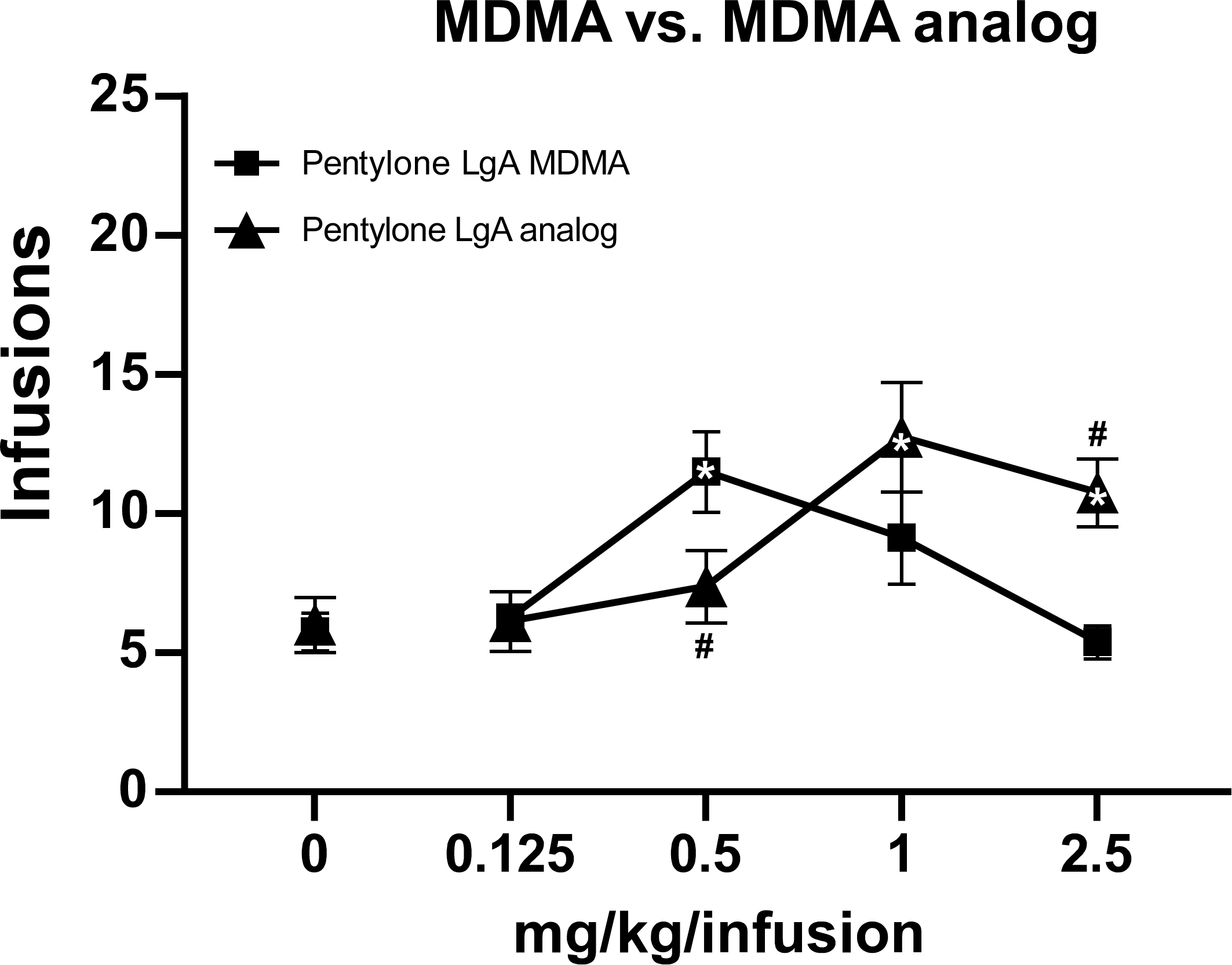

#### Figure S Self-administration of MDMA or MDMA-analog in female rats trained to self-administer Pentylone.

*Mean (±SEM) infusions of MDMA or MDMA-analog obtained by Pentylone self-administration trained rats (N=8). Mixed effects analysis confirmed a significant main effect of Dose [F(4,28)=5.857; P = 0.0015] and of the Dose X Group Interaction [F(4,28)=6.603; P = 0.0007]. The post hoc test confirmed the number of MDMA infusions (0.5 mg/kg/infusion) were significantly higher compared to vehicle, whereas the number of MDMA analog infusions was higher at 1 and 2.5 mg/kg/infusion doses. A significant difference from saline, within group, is indicated with * and a significant difference between groups is indicated with #.*
